# Metabolically intact nuclei are fluidized by the activity of the chromatin remodeling motor BRG1

**DOI:** 10.1101/2024.04.12.589275

**Authors:** Fitzroy J. Byfield, Behnaz Eftekhari, Kaeli Kaymak-Loveless, Kalpana Mandal, David Li, Rebecca G. Wells, Wenjun Chen, Jasna Brujic, Guilia Bergamaschi, Gijs J.L. Wuite, Alison E. Patteson, Paul A. Janmey

## Abstract

The structure and dynamics of the cell nucleus regulate nearly every facet of the cell. Changes in nuclear shape limit cell motility and gene expression. Although the nucleus is generally seen as the stiffest organelle in the cell, cells can nevertheless deform the nucleus to large strains by small mechanical stresses. Here, we show that the mechanical response of the cell nucleus exhibits active fluidization that is driven by the BRG 1 motor of the SWI/SNF/BAF chromatin-remodeling complex. Atomic force microscopy measurements show that the nucleus alters stiffness in response to the cell substrate stiffness, which is retained after the nucleus is isolated and that the work of nuclear compression is mostly dissipated rather than elastically stored. Inhibiting BRG 1 stiffens the nucleus and eliminates dissipation and nuclear remodeling both in isolated nuclei and in intact cells. These findings demonstrate a novel link between nuclear motor activity and global nuclear mechanics.

## Introduction

The nucleus is generally considered the stiffest organelle of the cell ^1–5^, with its deformation being the limiting factor in the ability of cells to squeeze through tight spaces of the crowded cellular environment. Measurements of nuclear stiffness, usually by atomic force microscopy or micro aspiration, show that apparent nuclear stiffness is weakly dependent on deformation rate and increases with increasing force, and provide values on the order of 5 to 10 kilopascal for the Young’s modulus ^6^ ^7^ ^8^, which is significantly greater than the stiffness of the cell’s actin-based cortex and orders of magnitude stiffer than whole suspended cells and the perinuclear or cytoplasmic volume of the cell, commonly reported to have elastic moduli on the order of 100 Pa ^9–11^. This model of a rigid nucleus sitting within a soft cell interior is inconsistent with the fact that cells can deform the spherical nucleus to a highly elongated shape when a cell is embedded within soft polymer networks such as those formed by a few milligrams per milliliter of collagen ^12^. The contractile stress generated by actomyosin in a non-muscle cell is generally on the order of a few kilopascals at most, and even if the contractile stress were higher, the soft collagen matrix surrounding the cell, a few 100 Pa for 3 mg/ml collagen, could not support much greater stresses, even if the collagen stiffens under deformation. A perhaps more direct demonstration that the nucleus might not be as stiff as often assumed, is the finding that lipid droplets within primary human hepatocytes that exert less than 200 Pa contractile stress are able to deform the nucleus, while retaining their spherical shape ^13^ ^14^. The high surface tension of the lipid droplet can plausibly account for the rigidity of the droplet, but the low contractile force characteristic of hepatocytes cannot account for the large local strain at the droplet-nucleus interface if the nucleus is rigid.

Most studies of nuclear stiffness are made either in intact cells, where the perinuclear intermediate filament network or the overlying actin cortical network are likely to contribute to the elastic resistance even when the measurements are made directly over the nucleus, and accurately accounting for the contributions of these multi-layered elastic materials is difficult. Alternatively, measurements of nuclei purified from cells after the cell’s plasma membrane has been disrupted and the nuclei separated from the cell remnants by centrifugation are confounded by the fact that these nuclei have lost soluble contents by diffusion through the nuclear pores and are no longer osmotically balanced with the cytoplasm. To overcome these limitations, in this study nuclei have been prepared by centrifugation of cells attached to a glass or hydrogel surface by a mechanism in which the nucleus, which is denser than the rest of the cell, emerges from the cell body with an intact plasma membrane wrapped around it, but no cytoskeleton, ribosomes, endoplasmic reticulum, or other organelles. The space between the intact nucleus and the plasma membrane contains approximately 500 nanometer thickness of cytoplasm containing glycolytic enzymes capable of generating ATP. This isolated, metabolically intact nucleus, termed a karyoplast, has been studied biochemically and morphologically ^15–17^, but not physically.

In this study isolated karyoplasts are characterized morphologically and deformed by atomic force microscopy using a flat cantilever to probe the full nucleus, over a wide range of deformation strains, frequencies, and time scales. Both the increased resistant force as the nucleus is deformed and the decrease in resistance as external forces are removed reveal the contributions of both viscous dissipation and elastic storage on the response of nuclei to mechanical stress. The effects of disrupting nuclear lamins, chromatin, cytoskeletal proteins, osmotic pressure, glycolysis, and surrounding polyelectrolytes that mimic the cytoskeleton on karyoplast stiffness have been measured. The results show that the mechanical response of nuclei cannot be explained by a passive viscoelastic model but requires the activity of ATPases, consistent with the properties of an active material that can change shape in response to even small forces, if they are applied over a long time, while allowing for full recovery of the original nuclear shape when the force is removed. Inhibition of the ATPase motor RNA polymerase II has little effect on karyoplast rheology, but inhibition of the ATPase Brahma-related gene (BRG-1) stiffened the karyoplast and abrogated dissipation, mimicking the effect of global ATP depletion, suggesting that movement by the Brg/Brm-associated factor (BAF) chromatin remodeling complex is essential for the active deformation of the nucleus.

## Results

The method by which karyoplasts are prepared is shown schematically in Figure 1. Cells are cultured either on glass cover slips or hydrogel substrates for 24 hours before reaching confluence. The cover slip with or without a gel is then placed into a modified centrifuge tube containing a spacer, below which is another cover slip coated with polylysine (1B). Before centrifugation, the nucleus in a well spread cell is typically a highly elongated oblate ellipsoid. After centrifugation, the nucleus appears as a sphere, wrapped in a sealed plasma membrane that has separated from the rest of the cell, termed a cytoplast (1C).

**Figure 1.**
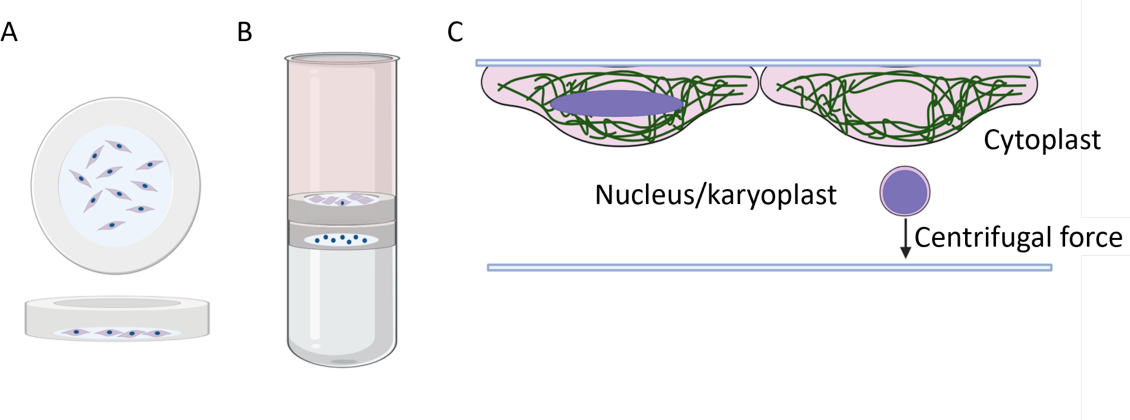
Nucleus isolation from mouse embryo fibroblasts using centrifugation. (A) Top and side view of a centrifuge tube insert with cultured mouse embryo fibroblasts. Inserts were made by bonding a glass coverslip to a ring of PDMS. (B) Ultracentrifuge tube containing enucleated cells in top insert and isolated nuclei collected on poly-d-lysine coated bottom insert. PDMS was cured in the base of the ultracentrifuge tube to stabilize inserts during centrifugation. (C) Side view of a cell before and during centrifugation, illustrating the isolation of a nucleus based on density. Notably, the isolated nucleus is surrounded by the donor cell’s plasma membrane and a thin layer of cytoplasm, also referred to as a karyoplast.

Figure 2 shows images of cells and their nuclei after the centrifugation process, where cells are stained for tubulin, vimentin, and chromatin. Enucleation of cells was performed using either normal mouse embryo fibroblasts or fibroblasts derived from a vimentin null mouse. Not all of the wild type cells have lost their nuclei under the relatively gentle conditions of this experiment, but almost all of the cells lacking vimentin have become enucleated (Fig. 2A). The efficiency of enucleation is shown quantitatively in Figure 2B. At a constant time of centrifugation (50 min), the percentage of cells that becoming nucleated is strongly dependent on the centrifugal force and is significantly more efficient in vimentin-null cells compared to wild type fibroblasts, reaching efficiencies above 80% with a few 1000 * g centrifugal force. At a constant centrifugal force, the efficiency of nucleation is strongly time dependent and again more efficient for vimentin null than wild type cells. A large fraction of nuclei can be pulled out of a cell within 50 minutes after the centrifugation has started. The appearance of the karyoplasts after their separation from the cells is shown in Figure 2C. In contrast to the flattened nucleus in the adherent cell, the karyoplasts are spherical. Figure 2D shows that the volumes of the spherical nuclei are indistinguishable between wild type and vimentin null cells and are similar to the values of the nuclei measured by light sheet microscopy in intact cells of the same cell types ^18^. The large variance of karyoplast volumes parallels the variance of whole cell volumes ^18,19^ and suggests that there is not a strong selection bias to remove either small or large nuclei, consistent with the ability to remove nuclei from nearly all of the cells, at least for vimentin null fibroblasts.

**Figure 2.**
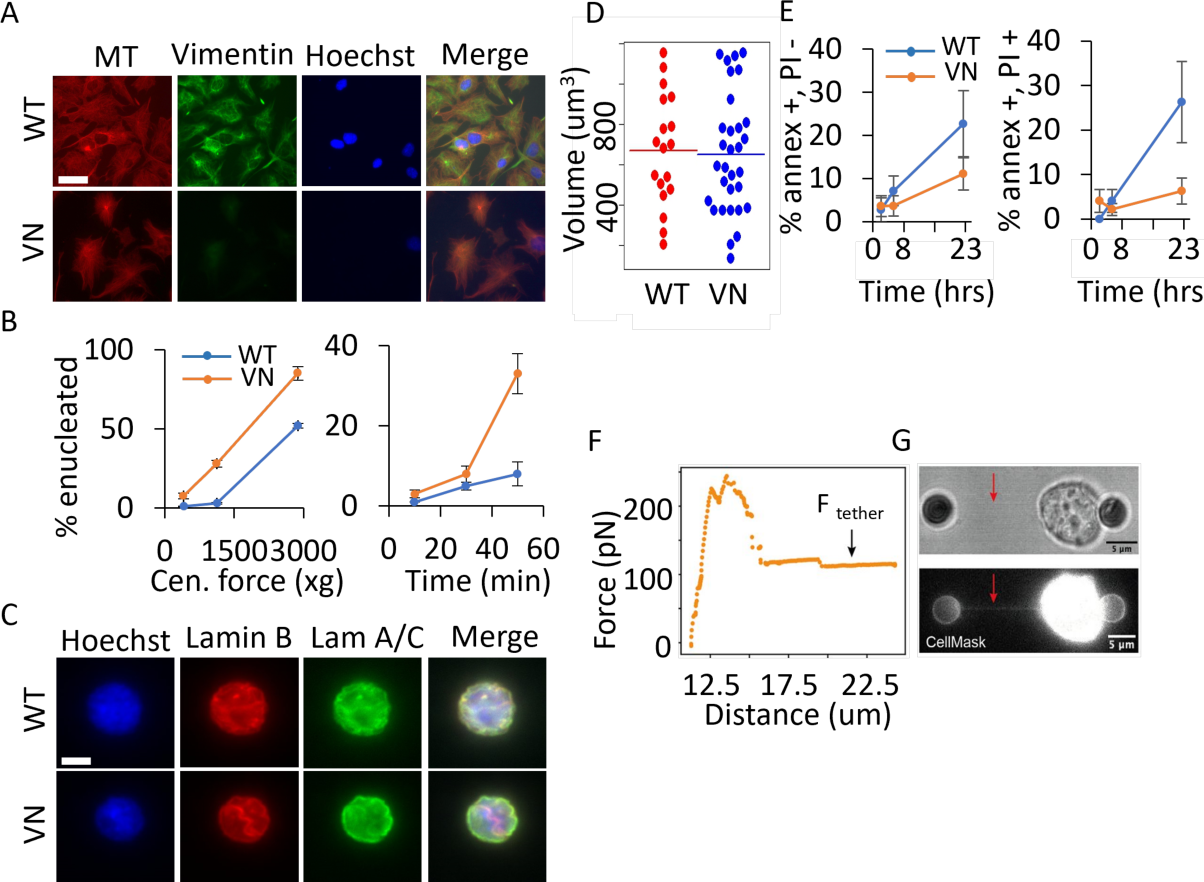
(A) Fluorescence images of wild type and vimentin null mouse embryo fibroblasts after enucleation. Scale bar = 25 um. (B) Analysis of enucleation rates with increasing centrifugal force for 50 mins at a centrifugal force of 2880 x g from 10 to 50 mins. (C) Fluorescence images of chromatin and lamins in nuclei isolated from wild type and vim null fibroblasts. (D) Scatter plot showing the volume of nuclei isolated from wild type and vimentin null fibroblasts. (E) Analysis of PS exposure and propidium iodide entry into isolated nuclei up to 22 hours after isolation. (F) Force vs distance pulled as a membrane tether is pulled from the surface of a karyoplast. (G) Bright field and fluorescence image of tether pulled from a karyoplast with fluorophore-labeled plasma membrane lipid bilayer,

The integrity of the plasma membrane is largely maintained in the karyoplasts after centrifugation, as shown in Figures 2E and F. Only a small fraction of the karyoplasts stain positive for annexin V, a marker that reveals the presence of phosphatidyl serine on the external leaflet of the plasma membrane, a sign that the cell or karyoplast has either undergone membrane disruption or has lost metabolic activity and the ability to maintain the enzymatic activity responsible for lipid bilayer asymmetry. We also find only a small fraction of propidium iodide-positive karyoplasts, which indicates disrupted plasma membranes, allowing the dye to enter the nucleus and stain the chromatin. Figure 2F shows that karyoplasts remain largely annexin V negative for many hours after their isolation, especially for those removed from vimentin null cells. 90% of the karyoplasts prepared from these cells maintain plasma membrane integrity and asymmetry for at least 23 hours after their isolation, during which time the rheologic measurements of this study are done. The plasma membrane also has excess surface area beyond the minimum to surround the spherical nucleus, because membrane tethers can be pulled from it with small forces similar to those required to pull tethers from an intact cell. To demonstrate this, we performed dual-trap optical tweezers experiments using two polylysine-coated beads to manipulate karyoplasts. This method facilitates localized force application directly onto the karyoplast membrane and allows precise control over bead attachment. When pulling on karyoplasts, we observed a distinct drop in the force data (Figure 2F) followed by a plateau F_tether_, a characteristic signature observed during tether pulling from cells ^20,21^. Simultaneously, fluorescence imaging of the karyoplast membrane, achieved by staining the karyoplasts with CellMask enabled real-time visualization of membrane stretching. Interestingly, we observed fluorescence along the tethers being pulled (Figure 2G), indicating that their origin comes from the plasma membrane of the karyoplasts.

The rheologic response of karyoplasts is measured by atomic force microscopy (AFM) as shown in Figure 3. To perform AFM studies, we use a large tipless silicon nitride cantilever that can make wide contact across the upper surface area of the karyoplast. The AFM cantilever is lowered on to the nucleus, and the resulting force as the entire nucleus is deformed is plotted against the change in nuclear height, as shown in Figure 3B. Similar to the finding that the appearance and size of nuclei are similar when prepared from wild type or vimentin null cells, the apparent stiffness is also similar. An apparent Young’s modulus can be calculated from appropriate fitting to the force deformation curves (Methods) and reveals an apparent Young’s modulus between five and six kilopascals (kPa) for both wild type and vimentin null cells.

**Figure 3.**
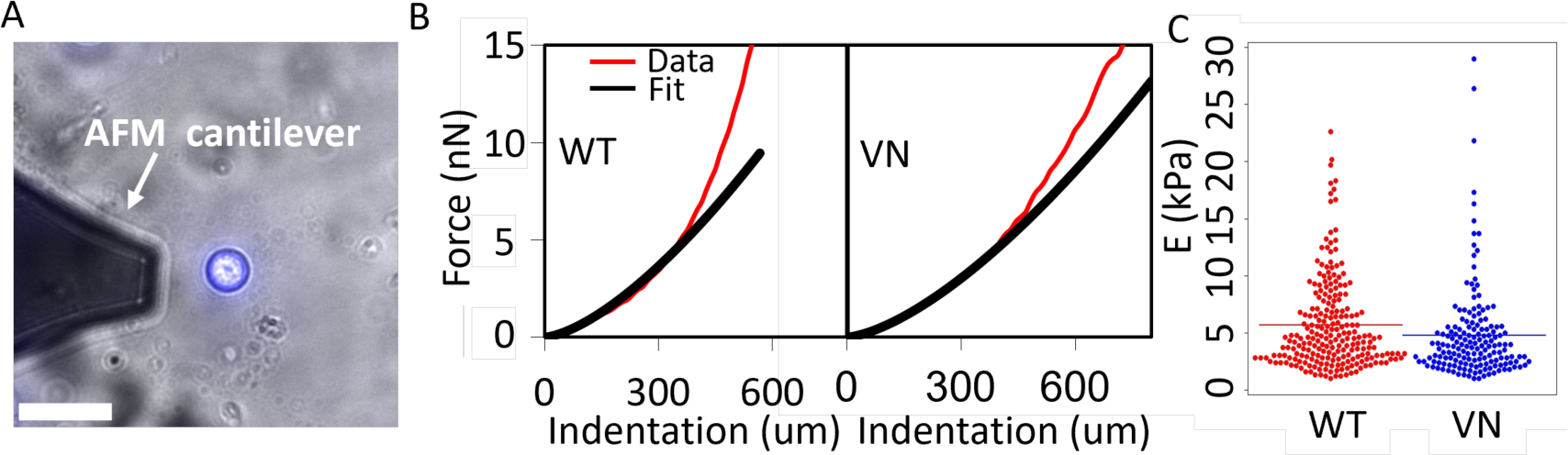
Mechanical properties of nuclei isolated from wild type and vimentin null mouse embryo fibroblasts. (A) AFM cantilever positioned next to a karyoplast attached to a poly-D-lysine coated surface. Scale bar = 20 um. (B) Force versus indentation curves for karyoplasts isolated from wild type and vimentin null fibroblasts fit to the Hertz model. (C) Scatter plot of the Young’s moduli of nuclei isolated from each cell type. p = 0.02

To determine which structures within the nucleus are responsible for its mechanical properties, karyoplasts have been treated with various agents to disrupt their membranes, the nuclear lamina, or chromatin. Since the plasma membrane is impermeable to large solutes, it was locally destabilized using digitonin, which clusters cholesterol and opens pores in the plasma membrane bilayer, but has no discernable effect on the nuclear membrane, which does not contain cholesterol. Permeabilization of the plasma membrane by digitonin has little effect on the overall nuclear morphology as shown in Figure 4A and B, although there appeared to be effects on the intensity and distribution of Hoechst staining. However, when DNase1 was added to the medium containing the karyoplasts and allowed to diffuse within the nucleus there was an abrupt change in nuclear size and stiffness.

**Figure 4.**
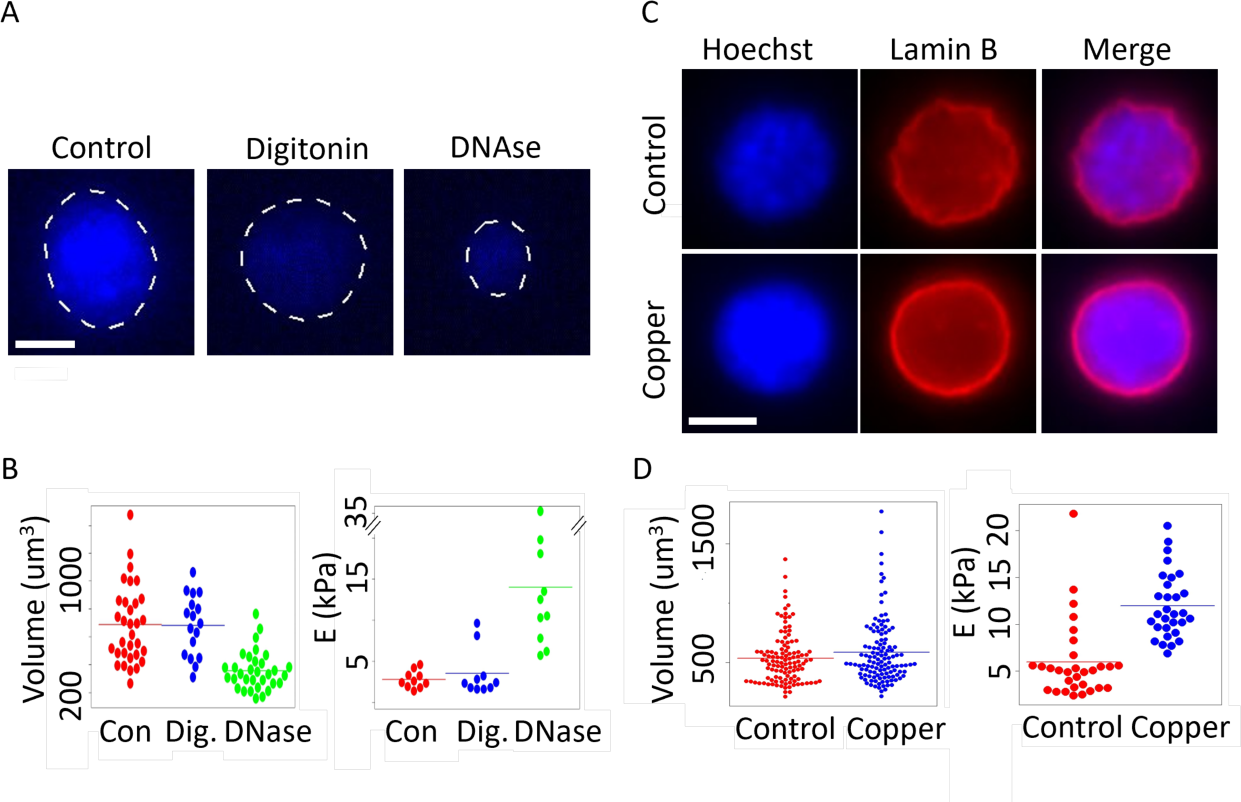
Figure 4. Volume and mechanical properties of isolated nuclei treated with DNase or copper. (A) Hoechst staining of isolated nuclei under control, digitonin, and digitonin + DNase treatments). (B) Scatter plots of volume (digitonin vs DNase, p =0.00001) and Young’s moduli (Digitonin vs DNase, P = 0.005) of nuclei corresponding to the treatments in panel (A). (C) Fluorescence images of nuclei treated with 1 mM copper sulfate. (D) Scatter plots representing the volume (p = 0.12) and Young’s moduli (p = 0.0000001) of nuclei corresponding to the treatments in panel (C). Scale bar = 5 µm.

The loss of Hoechst staining as shown in Figure 4A confirms that much of the DNA was removed from the karyoplast, and this perturbation led to a change in nuclear volume, dropping from approximately 700 to 250 µm^3^. Along with the change in volume, the remaining karyoplasts became much stiffer, increasing in apparent Young’s modulus to nearly 15 kPa. Perturbing the chromatin by condensing the DNA with copper had a very different effect on nuclear morphology and stiffness. As Figures 4C and D show, copper caused the DNA staining to become somewhat more clustered in the center of the karyoplast and increased apparent staining of lamin B, presumably because the condensation of chromatin allowed better access of the anti lamin B antibody to the nuclear lamina. These perturbations did not change the nuclear volume but approximately doubled the stiffness of the nuclei.

Next, we examined the role of the nuclear lamina on karyoplast stiffness. Figure 5A shows nuclei before and after treatment with CDK 1, a kinase that phosphorylates nuclear lamins and leads to their disassembly from the nuclear membrane. The loss of lamin staining after CDK1 is quantified in figure 5B. Despite loss of the lamin network, the volume of the karyoplasts was unchanged (Figure 5C). In contrast to the large effect of destabilizing nuclear lamins on local high curvature deformations of the nuclear membrane or when nuclei are stretched as cell traverse narrow spaces ^22–24^, Figure 5D shows that the apparent Young’s modulus decreases by less than 50% when the nuclear lamina is disrupted, consistent with the hypothesis that internal structures within the nucleus dominate this global response, at least when the nucleus is initially spherical.

**Figure 5.**
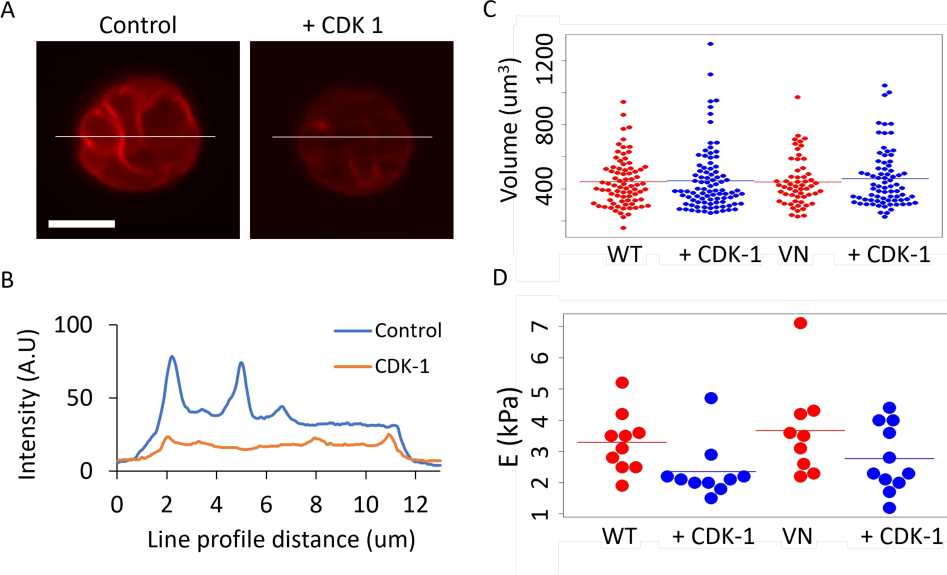
Volume and mechanical properties of nuclei treated with cyclin dependent kinase-1 (CDK-1). (A) Lamin B staining of isolated nuclei treated with CDK-1. Bar = 10 um. (B) Line profiles show decreased fluorescence intensity of lamin B staining after treatment with CDK-1. (C) Scatter plot of the volume of nuclei isolated from mEF WT and mEF VN with and without CDK-1. (D) Scatter plot of the Young’s moduli of nuclei isolated from mEF WT and mEF VN with and without CDK-1 (WT vs WT+ CDK-1, p = 0.04).

Similarly, treatment with cytochalasin, an inhibitor of cellular actin polymerization or azide, an inhibitor of ATP generation by mitochondria, had negligible effect on nuclear stiffness or morphology. These effects are expected since the cytoplasmic cytoskeleton and mitochondria are essentially absent from the plasma membrane-coated nuclei that constitute karyoplasts (Supplemental Figure 1A).

In contrast to the relatively small effects of inhibiting the nuclear lamina or the cytoskeleton, osmotic stresses on the isolated nuclei had large effects on both the volume and stiffness of karyoplasts. Figure 6A shows how the volume of karyoplasts is altered under hyperosmotic pressure caused by increasing the salt concentration in the surrounding medium, or by hypotonic media. The volume of the karyoplast decreases by less than 50% under an osmotic stress generated by 300 NaCl, corresponding to > 100 kPa, and the volume increases by nearly a factor of 3 when the osmotic pressure of the medium is decreased by removal of 140 mM NaCl. Along with these changes in nuclear volume, both hyper- and hypo-osmotic stresses had a large effect on the apparent Young’s modulus of the nuclei as shown in Figure 6B (10% PBS vs PBS, p = 0.0009; PBS vs +300 mM NaCl, p = 0.02). Hypoosmotic swelling, which led to a nearly threefold change in volume also led to approximately a threefold decrease in elastic modulus. In contrast, the relatively small change in volume under hyper osmotic conditions led to a seven times increase in elastic modulus, consistent with the idea that the nucleus can accommodate relatively large volume changes by increasing water content but cannot decrease volume by the same amount because of the large solid content held within the nuclear membrane even if it remains mostly water.

**Figure 6.**
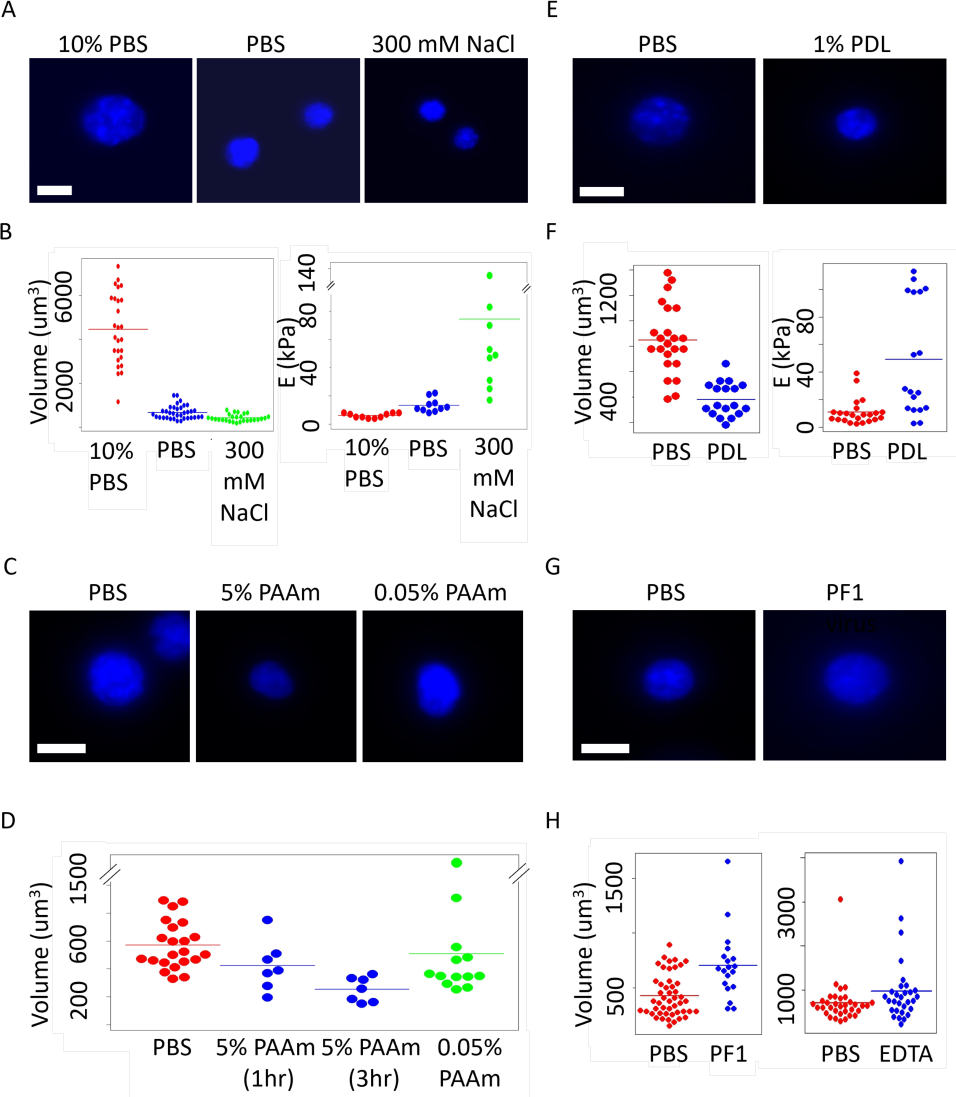
(A) Hoechst staining of karyoplasts in hypo (10% PBS + 90% dd H2O) and hyperosmotic (PBS+300 mM NaCl) solutions for 1 hr) (B) Scatter plots of volume and Young’s moduli of karyoplasts in hypo and hyperosmotic solutions. (C) Hoechst staining of isolated nuclei placed in 1% poly-D-lysine solution for 1 hr. (D) Scatter plots showing the volume and Young’s moduli of karyoplasts incubated with poly-D-lysine. (E) Hoechst staining of isolated nuclei in linear polyacrylamide (PAAm). (F) Volume analysis of nuclei incubated with linear PAAm. (G) Hoechst staining of nuclei incubated with 50 mg/ml PF1 virus for 4 hrs. Bar = 10 um (H) Volume of isolated nuclei suspended in PF1 virus and nuclei treated with 5mM EDTA. For all comparison p<0.05 except for PBS vs 0.095% PAAm.

Altering osmolarity by adding polymers to the outside medium had different effects on nuclear volume and stiffness than did changes in osmotic pressure induced by changes in salt concentration, and these effects depended on the electrostatic charge of the macromolecules. Figure 6C shows the effect of 5% polyacrylamide, an uncharged hydrophilic polymer, on karyoplast volume. This amount of macromolecule decreases the volume by approximately 20% after one hour and nearly 50% after three hours, but the loss of volume is nearly completely restored by dilution of the surrounding medium to remaining concentration of 0.05% (Figure 6D). Electrostatically charged macromolecules have significantly different effects on nuclear volume than do uncharged polymers like polyacrylamide. Figures 6 E,F shows that 1% of the cationic polyelectrolyte polylysine of molecular weight in the range of 70-150 kilodaltons, corresponding to a nominal osmotic pressure of less than 200 Pa decreased the nuclear volume by more than a factor of 2 and increased the apparent Young’s modulus by nearly 5x (PBS vs +PDL, p = 2 x 10 ^-8^). In contrast to the expected hyperosmotic effect of adding macromolecules to the surrounding medium, addition of an even larger weight concentration of the anionic polyelectrolyte filamentous bacteriophage PF1 led to a significant increase in nuclear volume (Figure 6 G,H). The mechanism by which addition of anionic polyelectrolytes increases nuclear volume, rather than decreasing it as would be expected from the osmotic pressure provided by the increased polymer solute concentration is not obvious but might be relevant to the setting in vivo where the nucleus is surrounded by a perinuclear cage formed of vimentin filaments with similar concentration (∼30 mg/ml) ^25^. In addition to more subtle polyelectrolyte effects, one possible effect of placing a large concentration of filamentous polyanions next to the nucleus is sequestration of divalent cations and their release from condensed chromatin. This possibility is supported by the finding that chelation of calcium by EDTA increases the volume of the karyoplast, although to a smaller extent than the filamentous anionic virus (Figure 6H). Parallel experiments with vimentin or actin filaments could not be done because such high concentrations of these biopolymers cannot be achieved in vitro to mimic the high concentration in which they are present in vivo. Measurements of nuclear stiffness could not be measured by AFM because of the viscoelastic resistance caused by the high concentrations the biopolymers added to the surrounding medium.

The apparent Young’s moduli of nuclei and cells measured by AFM is commonly calculated from the increasing force applied to the AFM cantilever as it is pressed into the cell or the karyoplast. Fitting these force-indentation curves to a Hertz relation assumes a linear elastic continuous material and yields an apparent Young’s modulus, but even an excellent fit of the Hertz relation to the force extension data does not guarantee that the cell or the nucleus is a linear elastic object. Figure 7A shows that when the AFM cantilever is first pushed into the nucleus and then pulled back there is a large difference in the force-displacement curves during indentation and recovery. This difference does not result from any damage or permanent change to the nucleus since repeated measurements of indentation and relaxation yield similar force-displacement curves. The repeatability of multiple deformations together with the extensive dissipation suggests an elastic structure acting in parallel with the highly dissipative material. As the height of the karyoplast decreases during compression its projected area increases, again nearly identically, during each compression cycle (Figure 7B) confirming that volume is conserved during these large deformations. At maximum, the compressive stress is near 50%, but the area does not reach the value seen for the nuclei in intact cells ^18^, showing that the cell on a rigid surface compresses its own nucleus more than can be done with the AFM at 50% strain. The reversibility of the indentation and the lack of permanent change is confirmed by the nearly perfect superposition of a decrease in nuclear height and an increase in the projected area (Figure 7C) consistent with a volume conserving deformation that can be repeated with very little change in the relation between nuclear height and projected area. In contrast to the simultaneous change in nuclear height and area, there is a significant lag between the vertical compression of the nucleus and the development of force pushing on the AFM cantilever. Figure 7D shows that the area increases more rapidly than the development of force during the compression phase but also continues to decrease even after the resisting force has relaxed nearly to the baseline. This lag between deformation and force, consistent with a highly dissipative material, is nevertheless reversible when the deformation is done again a few seconds after an initial indentation and recovery have occurred. Figure 7D shows that the hysteresis in the force-displacement curves is not the result of a simple viscous element because even though the dissipation is very large it is only very weakly rate dependent over nearly three orders of magnitude, and the measured Young’s modulus is also nearly independent of frequency.

**Figure 7.**
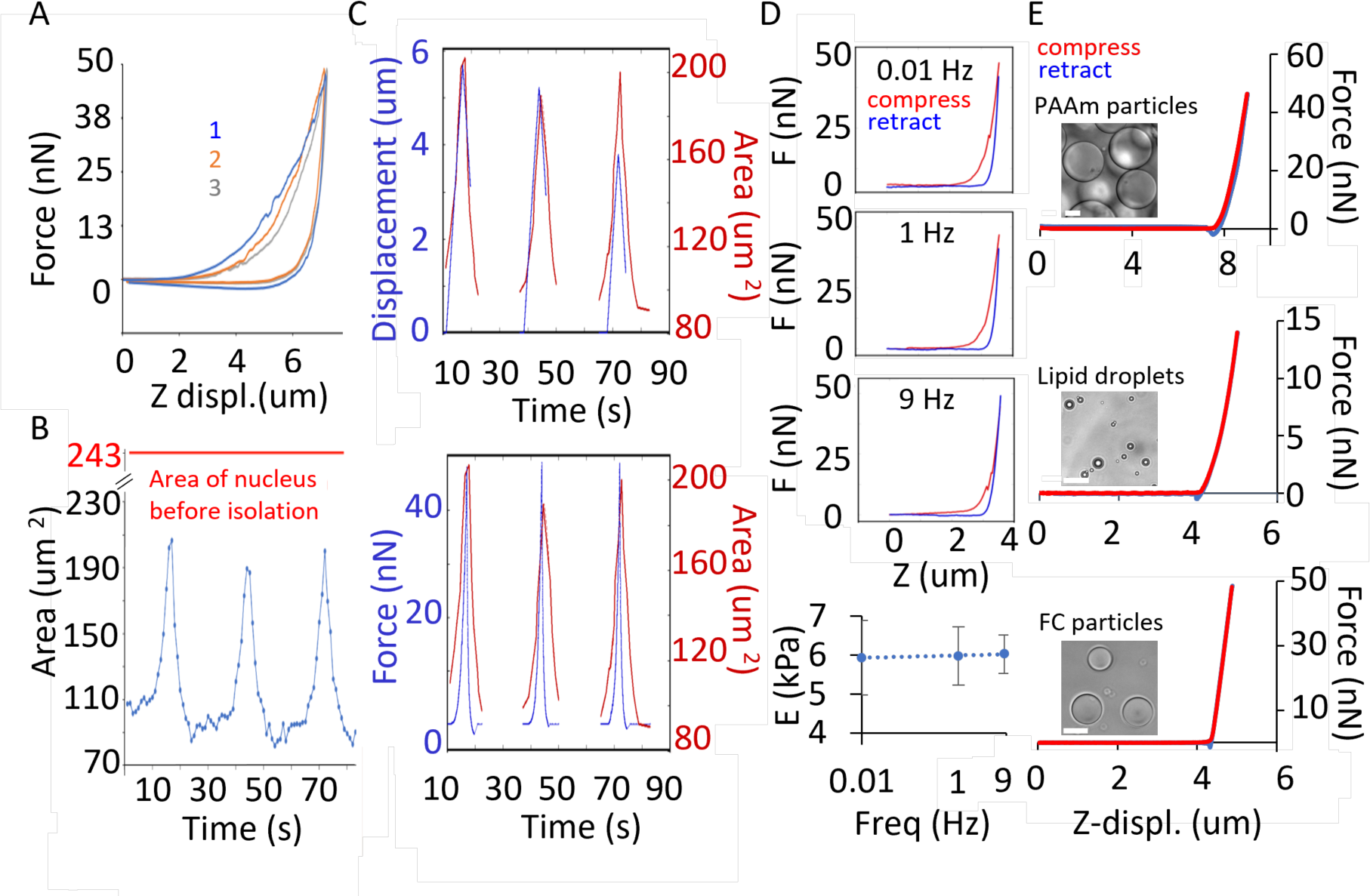
Comparison of dissipation during compression of nuclei and spherical microparticles. (A) AFM force curves showing consecutive compressions of a karyoplast at 0.1 Hz. (B) Area analysis of nucleus imaged during consecutive compressions. (C) Comparison of nuclear area change, nuclear deformation and applied force over time for three consecutive compressions of a karyoplast. (D) AFM compression curves for a nucleus compressed at rates of 0.01, 1 and 9 Hz and plot of Young’s modulus vs frequency (E) AFM compression of PAAm microparticles, lipid droplets and fluorocarbon particles compressed at 1 Hz. Bar = 20 um.

The possibility that the hysteresis between indentation and recovery might be an artifact of the AFM measurement, possibly due to unrecognized adhesion of the karyoplast to the substrate or the cantilever is countered by similar experiments done on three different kinds of small spherical objects. Figure 7E shows that the force indentation of a polyacrylamide sphere made by microemulsion is nearly purely elastic, as expected from the simple elastomeric cross-linked polyacrylamide gel. Figure 7E shows similarly that liquid droplets purified from liver or formed by fluorocarbon oil in which the elastic resistance comes entirely from surface tension that cannot relax during deformation show a nearly perfect correspondence between indentation and recovery during an AFM measurement similar to that applied to the nucleus.

The elastic modulus of the nucleus is not simply related to nuclear volume nor to the amount of chromatin, as shown in Figure 8. Karyoplasts were isolated from cells cultured on glass or polyacrylamide substrates with elastic moduli of 22 kPa or 5 kPa, which mimic the mechanical properties of soft tissues in which fibroblasts occur in vivo. The structures of cells, cytoplasts, and karyoplasts prepared on the three different substrates are shown in Figure 8A and confirm that nuclei can be removed from cells on each of the mechanically distinct substrates. After their removal from the cells, the karyoplasts look approximately similar, independent of the substrate stiffness (Figure 8B). Even hours after isolation of the nuclei from cells under these mechanical conditions, the volumes of the karyoplasts remain constant and indistinguishable from each other (Figure 8C), but the apparent elastic moduli of the nuclei are significantly different. Nuclei isolated from cells cultured on soft substrates had apparent Young’s moduli less than 30% of those isolated from cells cultured on glass (Figure 8D), even though the volumes of these nuclei and presumably the amount of chromatin within them is the same (Figure 8C). The hysteresis between compression and recovery is large in all cases and shows a trend toward more dissipation in nuclei isolated from cells on soft substrates (Figure 8E).

**Figure 8.**
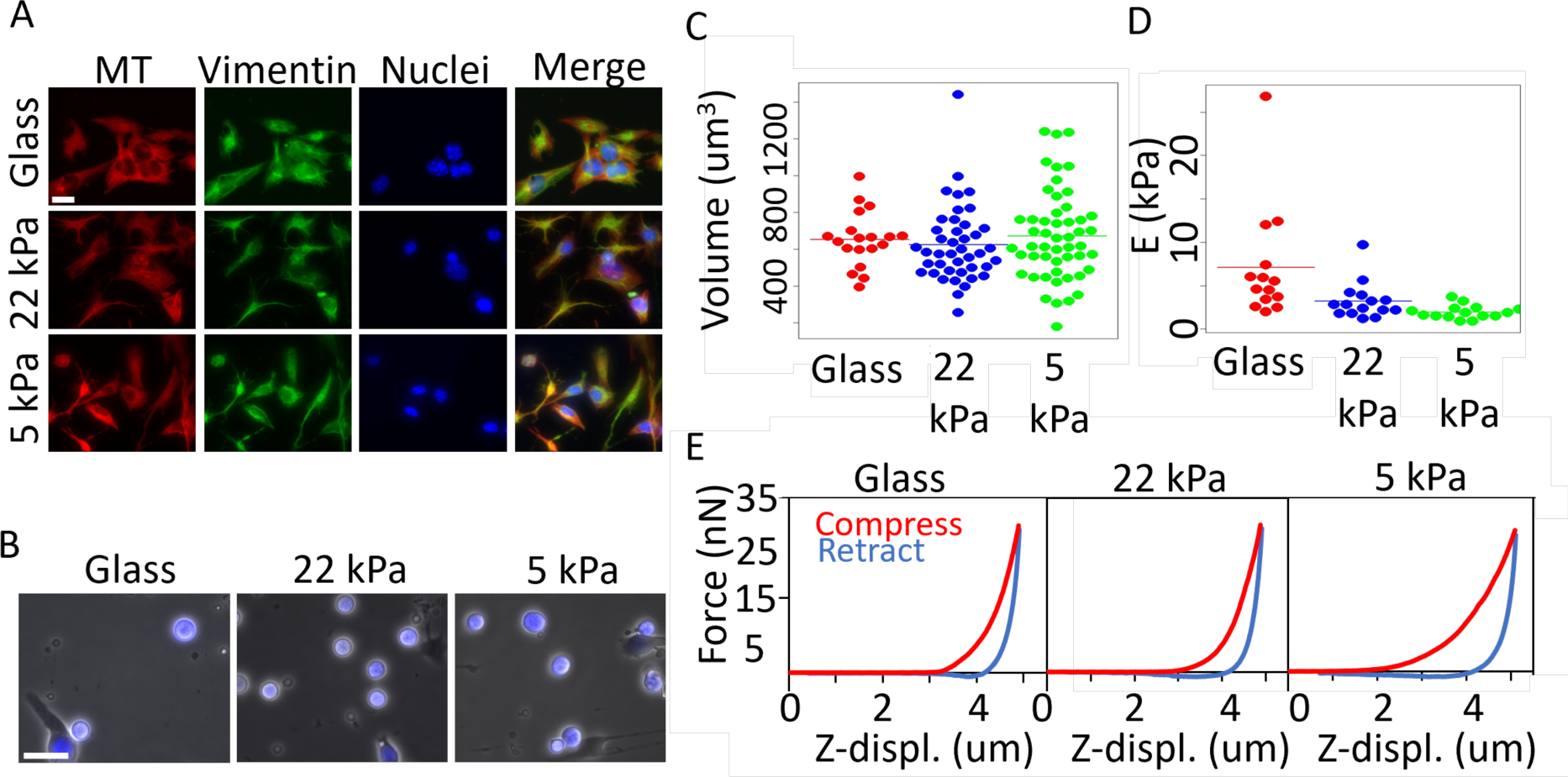
Characterization of nuclei isolated from cells grown on substrates of varying stiffness. (A) Fluorescence images of mEFs on collagen coated glass, 22 kPa PAAm and 5 kPa PAAm gels shown after the enucleation procedure. Bar = 20 um. Cells were cultured on each substrate for 24 hr before nucleus isolation. (B) Hoechst staining of nuclei isolated from cells cultured on glass, 22 kPa, and 5 kPa PAAm gels. Scatter plots of volume (C) and Young’s moduli (D) of nuclei isolated from cells grown on substrates of varying stiffness (Glass vs 22 kPa, p = 0.05; Glass vs 6 kPa, p = 0.01). (E) AFM force curves for nuclei isolated from cells grown on substrates of varying stiffness.

The very weak dependence of the apparent Young’s modulus on frequency and the high extent of rate independent dissipation shown in Figure 7 suggest that the mechanical dissipation occurring during deformation of the nucleus might result from an active process, rather than simple viscoelasticity. This possibility was tested by global inactivation of ATP-dependent motor proteins by treating the isolated karyoplasts with the glycolysis inhibitor 3 bromo pyruvate. Inhibiting ATP production from glycolysis has a large effect on both the elastic modulus and the amount of dissipation when a nucleus is compressed. Figure 9A shows that the hexokinase inhibitor 3-bromopyruvate produces an insignificant effect or possibly a small increase in the volume of karyoplasts, but figure 9B shows that it has a large effect on both stiffness and the amount of dissipation when nuclei are compressed. The apparent Young’s modulus increases by nearly a factor of 4 and the hysteresis between indentation and recovery is nearly eliminated.

**Figure 9.**
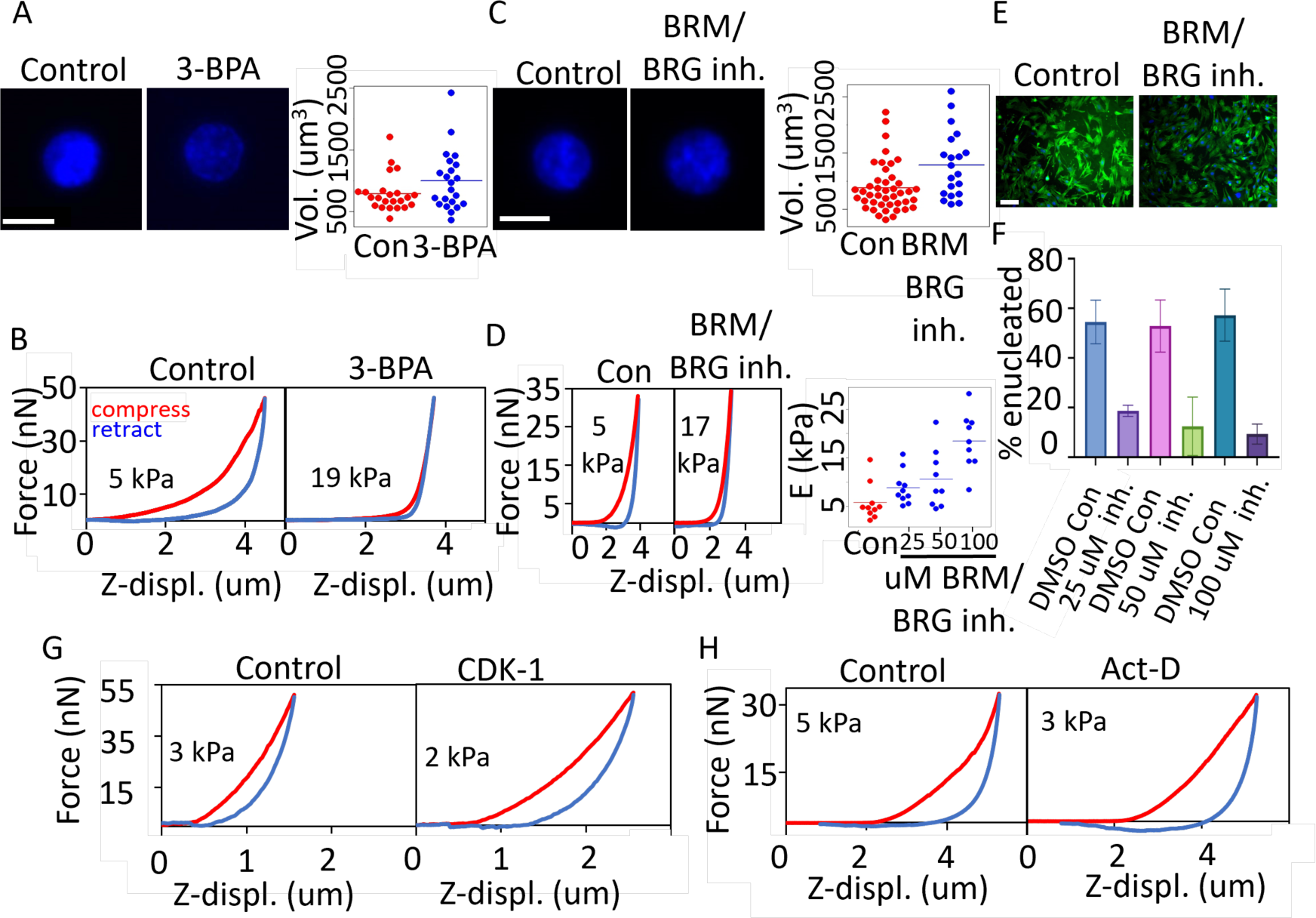
Characterization of isolated nuclei and cytoplasts after inhibition of ATP generation and BRM/BRG-1 ATPase activity. (A) Images and volume analysis of Hoechst-stained nuclei incubated with 100 uM 3-bromo-pyruvic acid for 2 hrs. (B) AFM compression curves of isolated nuclei treated with 3-bromo-pyruvic acid compared to DMSO control. (C) Images and volume analysis of isolated nuclei treated with 100 uM BRM/ BRG-1 inhibitor for 24 hrs (Control vs BRM/BRG-1 inhibitor, P-value = 0.009). (D) AFM compression curves for nuclei treated with 50 uM BRM/ BRG-1 inhibitor compared to DMSO control. (E) F-actin (green) and Hoechst (blue) staining of control and BRM/BRG-1 inhibitor treated mEF vim null cells. Images taken after enucleation procedure. (F) Analysis of enucleation efficiency after treatment with BRM/BRG-1 inhibitor. (G) AFM indentation-retraction curves for control and CDK-1 treated isolated nuclei. (H) AFM indentation-retraction curves for nuclei isolated from cells grown on substrates of varying stiffness. Scales bars: 10 µm A,C; 100 µm E.

The possibility that the dissipation is due to an active process driven by a chromatin remodeling enzyme or other motor protein was tested using specific inhibitors. The stiffening effect of ATP depletion was almost completely recapitulated by inhibition of a single ATPase in the nucleus, the BRG1 motor that drives chromatin reorganization by the BAF or SWI/SNF complex ^26^. Inhibiting BRG 1 slightly increases nuclear volume (Figure 9C), but it strongly increases the apparent Young’s modulus and decreases dissipation ((Figure 9D) in a concentration dependent manner, suggesting that the energy dissipation resulting from nuclear displacement is largely driven by active motions of chromatin.

To test whether mechanical effects of BRG1 inhibition can also alter nuclear rheology in the intact cell, adherent fibroblasts were treated with similar doses of the BRG1 inhibitor. After a few hours of incubation, the fibroblasts remained attached to the substrate and looked similar to those of the control uninhibited cells as shown in figure 9E. However, when subjected to centrifugal forces needed to pull the nuclei out of the cell to create a karyoplast, cells in which BRG1 was inhibited were unable to undergo this morphological transformation. Figure 9F shows that under conditions at which nearly 90% of control cells become enucleated, only 10% of nuclei could be removed from cells that had inactive BRG1.

Another perturbation of the karyoplast that stiffens it and eliminates dissipation is dissolution of chromatin by DNase I. Figure 4 shows how DNase I activity shrinks the nucleus and increases its Youngs modulus, and Supplemental Figure 1B shows that coincident with nuclear stiffening is loss of dissipation. Most other perturbations of the karyoplast led to either no change or an increase in the amount of dissipation, whether or not they produced altered the apparent Young’s modulus. Figure 9G,H shows that disassembly of the nuclear lamina, which makes the nuclei somewhat softer, led to a small increase in the hysteresis between indentation and recovery. Inhibition of RNA polymerase by actinomycin D had only a modest effect on nuclear volume (Supplemental Figure 1C) and a small softening effect on karyoplasts, with possibly an increase in dissipation (Figure 9 I,J). These results suggest that the energy dissipation is largely independent of the nuclear lamina but dependent on DNA and some element that actively promotes chromatin movement.

## Discussion

Formation of karyoplasts, in which an intact nucleus has been in continuous chemical equilibrium with the cytosol and in which large elastic structures of the cytoskeleton and cytoplasmic organelles have been removed while retaining an intact plasma membrane and continuous ATP production, permits mechanical investigations of nuclear structure and mechanics without confounding factors from the rest of the cell. Removal of nuclei from the cell by gentle centrifugation relaxes the stress that the cytoskeleton exerts on the nucleus and retains the chemical composition of the nucleus under variable physical conditions caused by such factors as substrate stiffness without the capacity to exchange nuclear contests with the rest of the cell. Our measurements of metabolically active nuclei within the karyoplast reveal some novel aspects of nuclear structure and mechanical properties. The major finding is that a large fraction of the work done to compress the nucleus is dissipated rather than elastically stored. This dissipation is not the result of viscoelasticity but rather results from active fluidization of the nucleus by the chromatin remodeling complex SWI/SNF, driven by the ATPase BRG1.

As reported earlier, nuclei from normal and vimentin null cells produce indistinguishable karyoplasts ^27^, but their formation is more efficient from cells lacking the perinuclear vimentin network. The volumes of karyoplasts are also indistinguishable from those of the intact cells from which they are prepared ^18^, suggesting that the osmotic balances that control nuclear size in the cell are maintained in the karyoplast. Once isolated the karyoplast volume is strongly dependent on osmotic conditions. Numerous recent studies have shown how nuclear and cell volumes are altered by osmotic pressures caused by changes in solvent concentration ^28–31^, and that nuclei become stiffer when they are osmotically compressed ^32^. Consistent with these studies, the results in Figure 6 show that nuclear volume can be increased strongly under hypoosmotic conditions but decreased only to a limited extent in hyperosmotic conditions, presumably because the solute content within the nucleus is near the point at which the activity of water is far from that in a dilute solution ^28^.

The osmotic sensitivity of nuclear volume is also seen when macromolecules, rather than small solutes, are the source of osmotic stress. In the case of polymers, the charge density on the macromolecule has a strong effect on the volume response of the karyoplast. Positively charged polymers are significantly more efficient than neutral polymers in creating a hyperosmotic environment in which the karyoplast shrinks. In striking contrast, a nominally hyperosmotic environment formed by a high concentration of anionic macromolecules, in this case the nanoscale filamentous FD virus, causes the nucleus to swell rather than shrink (Figure 6 G,H), as would have been expected due to the high concentration of polymer added to the medium. This swelling of the nucleus caused by anionic polymers is presumably due to polyelectrolyte effects that shift the balance of mobile counter ions between the anionic DNA inside the nucleus and the anionic macromolecule outside, but whether this effect is due only to sequestration of divalent cations by the external polyelectrolyte or to some other polyelectrolyte effect remains to be clarified. The swelling effect of PF1 virus on the nucleus might however account for the fact that nuclei within cells that have a dense perinuclear network of vimentin intermediate filaments, which have similar charge density as Pf1 virus ^33^, are larger than nuclei in cells without vimentin ^18^, even though when karyoplasts are formed from these cells therefore, they have the same volumes (Figure 2).

Numerous studies indicate that both the nuclear lamina and chromatin configuration play significant roles in shaping and maintaining the rigidity of the nucleus ^34^. Studies on isolated nuclei using pipette stretchers indicate that the level of euchromatin and heterochromatin mediates nuclear stiffness at relatively small strain, while at larger extensions lamin A/C control stiffening of the nucleus ^35^ with loss of lamin A/C decreasing stiffness by 36% at large strains. Measurements with micropipette aspiration also show an approximately 50% decrease in nuclear compliance with lamin A/C knockdown ^36^. Further, disruption of nuclear lamins increases nuclear deformations and can lead to nuclear abnormalities, such as blebbing and nuclear envelope rupture ^37^. Our results of AFM compression of isolated nuclei show similarly a decrease in nuclear stiffness upon treatment with CDK 1 that disassembles the nuclear lamina, and that the loss of lamina does not alter the degree of dissipation during nuclear compression. The relatively small effect of disrupting the nuclear lamina is also consistent with previous studies showing that deformation to strains below 40% are dominated by the nuclear interior with the lamina dominating at larger strains ^35^. In addition to the nuclear lamins, our study highlights a significant role of nuclear activity and the BRG1 motor of the SWI/SNF complex in mediating nuclear stiffness.

Perhaps the most striking rheological characteristics of karyoplasts are the high degree of dissipation when they are deformed to large extents, the reversibility when compressed to strains greater than 50%, and the dependence of mechanical dissipation on the generation of ATP and the activity of intranuclear motor proteins. The partly fluid nature of the nucleus has been previously shown, for example in striking images of nuclei being pulled into micro capillaries ^36^ and by measurements of chromatin deformation by a ferromagnetic bead in a magnetic field gradient, which also emphasized the fluidity of the nucleus ^38^. The ability of the nucleus to flow in a response to an external stress does not appear to be the result only of a high passive viscosity, but rather is due to active fluidization of the dense intranuclear core. This fluid nature of the karyoplast disappears when DNA is digested, glycolysis is prevented, or when the BRG 1 motor of the SWI/SNF complex is inhibited.

The high degree of karyoplast stiffening upon ATP depletion is closely paralleled by recent AFM studies of nuclear stiffness in intact cells before and after permeabilization. In the intact cell, the nuclear region has an apparent Young’s modulus of 5 kPa, but when the nucleus is exposed to the medium by detergent permeabilization of the cell and nuclear membranes, the elastic modulus rises to 22 kPa ^7^, very similar to the values shown in Figure 9. Other studies, however, report that the nucleus within the cell is stiffer than the insolated nucleus, because of cell-generated stresses that deform the nucleus and stiffen it by pre-tension ^8^.

The BRG1-containing chromatin remodeling complexes SWI/SNF or BAF have structural and dynamic properties consistent with their ability to fluidize the nucleus. These complexes can move chromatin or naked DNA at rates of 50 base pairs per second (bp/s) with a step size of 2 bp ^39,40^, implying that the turnover rate of the motor is >25 Hz, which is faster than the rates of deformation accessible to the AFM (Figure 7), and therefore consistent with the finding that although nuclear deformation is largely dissipative, it is not rate dependent over the range of our measurements. The force developed by this motor is also remarkably high, 12 nN ^40^, enabling it to make large movements in the dense environment of the nucleus.

The data presented here show that fluidization of the nucleus is eliminated when the BRG1 motor is inhibited, but other structures within the nucleus might also be important in determining both the Young’s modulus and the degree of dissipation. When karyoplasts are isolated from cells cultured on substrates with different stiffnesses, they retain very different rheological properties even long after their isolation from the cell. One well documented difference in nuclear structure that depends on substrate stiffness is the trafficking of the transcription regulatory proteins yap and taz into the nucleus. Whether alterations and transcription profiles are sufficient to change nuclear stiffness is not known. However, another structure that was recently shown to change strongly with substrate stiffness is the amount of polymerized nuclear actin ^41^. Networks of actin filaments form when cells are cultured on stiff substrates but only amorphous structures of similar concentrations of actin are seen in the nucleus of cells on soft substrates. The role of actin and myosin motors in the nucleus is much less understood than the function of these proteins in the cytoskeleton, but their presence in the nucleus is increasingly well documented, and our preliminary experiments show that 2,3-butanedione monoxime (BDM) which is a relatively nonspecific inhibitor of myosin affects karyoplast rheology, as shown in Supplemental Figure 1D. The possibility that actin filaments might contribute to karyoplast rheology is supported by earlier studies showing that the SWI/SNF complex contains a subunit of beta actin and actin related proteins and nucleates actin filaments ^42^. Moreover, when actin polymerizes within the nucleus on a stiff substrate it is bound to the SWI/SNF complex, and this binding is essential for the transcriptional activity of yap and taz regulators ^41^.

The current study has several limitations. The nuclei studied are only from fibroblasts, and only from cells in interphase, where the nucleus can be released intact by centrifugation. The data strongly suggest that the nucleus is not the stiffest organelle of the cell; internal membraneless organelles such as the nucleolus are stiffer ^43^ as are lipid droplets, even when their size is similar to that of the nucleus. Much remains to be learned about how nuclear composition relates to nuclear mechanics, and the method of making karyoplasts that retain much of the native structure and activities of the intact nucleus provides a useful tool for further studies.

## Materials and Methods

### Cell Culture

Wild-type mouse embryonic fibroblasts (mEF WT) and vimentin null fibroblasts (mEF vim null) were cultured in high glucose DMEM (Corning) plus 10% FBS (Sigma-Aldrich), 0.1 ug/ml Penicillin, 0.1 mg/ml Streptomycin (Mediatech), and 5% non-essential amino acids (Gibco). Cells were maintained at 37 °C in a 5% CO2 environment with saturated humidity.

### Centrifuge Tube Insert and Polyacrylamide Gel Fabrication

As previously described ^27^ inserts for thick-walled polycarbonate ultracentrifuge tubes (32 ml capacity, Beckman Coulter, Brea, CA) were prepared by bonding air plasma-treated round 18 mm diameter glass coverslips to polydimethylsiloxane (PDMS) rings (inner diameter:13mm, outer diameter:20 mm, height :3mm). Next, inserts were functionalized by treating with (3-aminopropyl)trimethoxysilane. Polyacrylamide (PAAm) gel mixtures with Young’s moduli ranging from 5 kPa (5% acrylamide, 0.15% bis-acrylamide) to 22 kPa (8% acrylamide, 0.48% bis-acrylamide) were then added and polymerized under a Rain-X treated glass coverslip. To prevent cell attachment to the uncovered glass surface, 2% agarose was added to the periphery of the gels. Inserts with PAAm gels were coated with 0.1 mg/ml collagen by first functionalizing the gel surface with sulfo-SANPAH and incubating with ligand overnight at 4 degrees. Inserts without PAAm gels underwent air-plasma treatment and were also incubated with ligand overnight. In some cases, fluorescent microspheres were added to the PAAm gels to enable traction field microscopy studies ^44^. Since the karyoplasts retain an intact plasma membrane, they adhere to the collagen-coated gel surface. Supplemental Figure 1E shows the traction field immediately below an adherent karyoplast. The small strain field is consistent with gel deformation cause by the adhesion energy of the spherical karyoplast, similar to that formed by adhesion of a phospholipid vesicle ^45^, and shows that the remaining cytoplasm does not spread on the surface of the hydrogel.

### Nucleus Isolation

Nuclei were isolated as previously described ^27^ based on the methods of ^46,47^ with some modifications. Cells were cultured on the prepared inserts for 24-48 hours before isolation of nuclei. Just before isolation, cells were incubated with 5 ug/ml Hoechst (diluted in cell culture media) for 30 mins. Subsequently, samples were centrifuged at speeds ranging from 405 to 2880 x g for up to 50 mins. The centrifugation was performed in DMEM supplemented with 1% FBS and 2 ug/ml cytochalasin D at 37 °C using a Beckman-Coulter Optima LE-80 K ultracentrifuge with an SW-28 rotor. To stabilize the inserts during centrifugation, PDMS was cured in the base of ultracentrifuge tubes to create a flat surface. During centrifugation, isolated nuclei were collected in inserts coated with 0.1 mg/ml poly-d-lysine. Enucleation efficiency was assessed by imaging cells after nucleus isolation and analyzing the percentage of cells lacking nuclei, as determined by the absence of Hoechst staining. Additionally, the volume of isolated nuclei was calculated assuming a spherical shape based on their radii.

### Fluorescent Staining of Isolated Nuclei and Cytoplasts

Freshly isolated nuclei were incubated with Annexin V and propidium iodide for up to 24 hrs to determine exposure of phosphatidylserine (PS) and integrity of the plasma membrane surrounding the nuclei, respectively, using an apoptosis detection kit (Life Technologies, V13241). For immunostaining, both nuclei and cytoplasts were fixed with 4% paraformaldehyde for 10 mins at room temperature. Samples were then permeabilized with 0.1 % Triton X-100 for 5 mins and incubated with 1 % bovine serum albumin (BSA) for 30 mins. For isolated nuclei, incubations were then made with lamin A/C (Cell Signaling Technologies, 4777S) and lamin B (ABCAM, ab16048) antibodies (diluted 1:500 in 1% BSA) for 1 hr. For cytoplasts, incubations were made with alpha tubulin monoclonal antibody (Bio Rad, MCA77G) and vimentin antibody (Novus Biologicals, NB300-223) also for 1 hr. Samples were then rinsed with 1% BSA and incubated for 1hr with the following fluorescently labeled secondary antibodies: goat anti-chicken IgY (Invitrogen, A11039), goat anti-mouse IgG (Invitrogen, A11029), and goat anti-rat IgG (Invitrogen, A11007). These antibodies were diluted 1:500 in 1% BSA. Samples were then rinsed with PBS and imaged using a 63x objective. Analysis of fluorescence intensity was performed using Image J.

### Treatments of Isolated nuclei

Hoechst-stained isolated nuclei, attached to glass coverslips unless indicated otherwise, were exposed to the following treatments: for DNA degradation experiments, nuclei were permeabilized with 1% digitonin and treated with 1mg/ml DNase in PBS for 1 hr. In treatments involving copper, nuclei were permeabilized with 0.1 % saponin then treated with 1 mM copper sulfate in tris-buffered saline (TBS) for 1hr. To investigate osmotic effects, nuclei were exposed to PBS, 50% PBS/ 50 % H2O, 90% PBS/ 10% H20, PBS + 100 mM NaCl, or PBS + 300 mM NaCl for 1 hr. For nuclear lamin disassembly experiments, nuclei were treated with 90 pmol cyclin dependent kinase-1 (CDK-1) in the presence of 1 mM ATP for 1 hr in suspension. Additionally, nuclei were treated with either cationic 1% poly-d-lysine or neutral 5% linear polyacrylamide (PAAm) in PBS for 1 hr. Alternatively, isolated nuclei were incubated with 50 mg/ml anionic PF1 virus (Asla Biotech, Riga LV, dialyzed against PBS) in suspension for 4 hrs. ATP production by glycolysis was inhibited by 3-bromopyruvate (Cayman Chemical 19068). RNA polymerase was inhibited by Actinomycin D (Invitrogen A7592). The BRG1 motor of the BAF complex was inhibited by the dual BRM and BRG1 inhibitor BRM-014 (Cayman Chemical 36138).

Following these treatments, nuclei were either imaged with a Leica DM IRE2 microscope using a 63x objective or compressed using a Bruker Nanowizard 4 atomic force microscope (AFM) mounted on a Leica DMI600 fluorescence microscope.

### Fabrication of Polyacrylamide (PAAm) Microparticles

Polyacrylamide (PAAm) microparticles were fabricated through inverse emulsion polymerization. For this purpose, the surfactant polysorbate 85 (Span 85, Sigma-Aldrich) was dissolved at a 2% (v/v) concentration in 200 ml of cyclohexane solvent to stabilize the spherical morphology of particles. Due to free radicals being required to initiate the polymerization of PAAm, oxygen, a free radical trap, was removed from the synthesis system by linking a nitrogen tank to the flask’s rubber stopper. During the degassing process, a 10 mL PAAm solution was prepared using acrylamide, bis-acrylamide, ammonium persulfate (APS, Sigma-Aldrich), and phosphate buffered saline (PBS). The final concentration of acrylamide, APS and bis-acrylamide were kept constant at 4%, 0.1% (v/v) and 0.2% (v/v), respectively. Then, N,N,Nʹ,Nʹ-tetramethylethylenediamine (TEMED, Thermo Fisher Sci.) was added to the PAAm solution to yield a final concentration of 0.1% (v/v). After 10 seconds of vortexing, the PAAm solution was added dropwise into the cyclohexane/Span 85 mixture. The stirring rate was enhanced (∼1000 RPM) to make microparticles of the desired size. Once the polymerization reaction was completed (∼1 h), stirring was stopped, and microparticles were allowed to settle for 30 minutes. The supernatant was removed, and the microbeads were washed twice with 100% ethanol and pelleted by 5 minutes centrifugation at 400g. The microbeads were then immersed overnight, in PBS, on a shaker.

### Fabrication of fluorocarbon droplets

A monodisperse droplet emulsion was synthesized according to a protocol adapted from ^48,49^, as outlined in ^50^. Briefly, an equal volume of dimethoxydimethysilane (98%, Sigma-Aldrich) and (3,3,3-trifluoropropyl)methyldimethoxysilane (97%, Gelest) was mixed together with deionized water at 5 vol%. The mixture was homogenized by vortexing for 5 min and shaking on a shaker (∼ 1000 rpm) for 6 h. Ammonia (27 vol% aqueous solution, Sigma-Aldrich) was then added at 1 vol% and mixed by gently tumbling the container, and the droplets were left to grow on a rotator for 24 h. 100 mM SDS solution was added to the mixture to reach a final SDS concentration of 1 mM to prevent further growth and coalescence. The droplets were then washed three times by centrifugation with 1 mM SDS to remove ammonia and reaction byproducts, and finally stored in a 5 mM SDS solution until further use.

### Animal Studies

All animal work was carried out in strict accordance with the recommendations in the Guide for the Care and Use of Laboratory Animals of the National Institutes of Health. Animal protocols were approved by the Institutional Animal Care and Use Committee of the University of Pennsylvania (protocol #804031). ob/ob mice were obtained from the Jackson Laboratories (strain #000632) and were housed in a temperature-controlled environment with appropriate enrichment, ad libitum feeding of standard rodent chow and water, and 12h light/dark cycles. Euthanasia was carried out by CO_2_ inhalation followed by exsanguination.

### Lipid Droplet Isolation

Lipid droplets were isolated from ob/ob mouse livers harvested at 12 weeks with a sucrose density gradient as described previously, with minor modifications ^51^. Briefly, fresh livers were minced and washed with 1X DPBS, then homogenized in a Dounce tissue grinder (Wheaton) with 55.5% (w/v) sucrose in TE buffer (10 mM Tris-HCl [pH 7.4], 1 mM EDTA [pH 8.0]). The homogenate was centrifuged at 1000 x *g* for 10 min at 4°C. 8 mL of the supernatant was placed into a 50 mL conical, and 6 mL of 40% (w/v) sucrose in TE buffer was carefully added on top, ensuring that the two phases remained separate. 6 mL of 25% sucrose in TE buffer, 4 mL of 10% sucrose in TE buffer, and 8 mL of TE buffer were then added in a similar fashion before centrifugation at 2000 x *g* at 4°C for 30 min. The top layer containing the lipid droplets was transferred using a glass pipette to a 2 mL microcentrifuge tube and washed in 1x DPBS. All solutions contained protease and phosphatase inhibitors, used at the concentrations recommended by the manufacturer (cOmplete Protease Inhibitor Cocktail and PhosSTOP, Sigma Aldrich).

### Atomic Force Microscopy (AFM)

AFM measurements were made with a Nanowizard 4 (Bruker) mounted on a Leica DMI 6000 B microscope. Before each experiment, a calibration was made by determining the slope of the deflection of the AFM cantilever on a petri dish to determine the deflection sensitivity. Isolated nuclei were distinguished from cells that detached during centrifugation by a near complete overlap between bright field images and Hoechst staining.

Nuclei, PAAm microparticles, fluorocarbon microparticles and lipid droplets were compressed using tipless silicon nitride cantilevers (NP-O10, Bruker, Camarillo, CA) with a nominal spring constant of 0.35 N/m at frequencies of 0.01, 1, and 9 Hz and a maximum applied force of approximately 30 nN. All measurements were made at room temperature in PBS with a minimum of 10 samples per condition.

AFM force curves were analyzed using the JPK analysis software and fit to a double contact hertz model for a sphere being compressed between two planes ^52^:

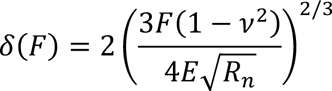

where δ is the deformation as a function of the force *F, v* is the Poisson’s ratio, *E* is the Young’s modulus and *R_n_* is the radius of the nucleus. In some cases the index of plasticity (IOP) was calculated as the relative difference in the areas below the indentation and retraction curves ^53^.

### Optical tweezers experiments

Before experiments, the microfluidic flow cell (Lumicks B.V) was passivated with 0.3% casein for about 1 hour, then rinsed with H2O and Nuclear Buffer (10 mM Hepes pH 7.5, 2mM MgCl2, 25mM KCl, 250 mM Sucrose,1mM DTT, 1mM PMSF). Carboxylated polystyrene microspheres (3% v/v, Spherotech) of 4.67 um diameter were washed (2000g for 1 min) and incubated with 0.05% poly-L-lysine (Sigma Aldrich) for 30 mins at RT on a rotating Eppendorf mixer; the final pellet was resuspended in 1mL Nuclear Buffer. Karyoplasts were stained with CellMask and finally diluted 1:4 in Nuclear Buffer (for a final volume of 150-200uL). For optical tweezer experiments, the karyoplast solution was flushed in channel 4, while the beads were flushed in channel 2. After catching two beads in the dual traps, the latter are moved to channel 4 and attached to a single karyoplasts; the karyoplast is then moved to channel 3 (containing Nuclear Buffer) where the stretching experiments are performed.

## Acknowledgements.

This work was supported by the US National Institutes of Health and the US National Science Foundation through grants NIH R35GM136259 and NSF CMMI-1548571 (PAJ) and NSF CMMI 2238600 and NIH R35GM142963 (AEP). We are also grateful for discussions with Qi Wen and Jay Yang, who suggested that the effect of anionic polyelectrolytes might relate to sequestration of divalent cations.

## Notes

### Competing Interest Statement

The authors have declared no competing interest.

## References

1 Cali, B. et al. Atypical CXCL12 signaling enhances neutrophil migration by modulating nuclear deformability. Sci Signal 15, eabk2552, doi:10.1126/scisignal.abk2552 (2022).

2 Agsu, G. et al. Reconstituting the Interaction Between Purified Nuclei and Microtubule Network. Methods Mol Biol 2430, 385–399, doi:10.1007/978-1-0716-1983-4_25 (2022).

3 Niethammer, P. Components and Mechanisms of Nuclear Mechanotransduction. Annu Rev Cell Dev Biol 37, 233–256, doi:10.1146/annurev-cellbio-120319-030049 (2021).

4 Miroshnikova, Y. A. & Wickstrom, S. A. Mechanical Forces in Nuclear Organization. Cold Spring Harb Perspect Biol 14, doi:10.1101/cshperspect.a039685 (2022).

5 Dahl, K. N., Ribeiro, A. J. & Lammerding, J. Nuclear shape, mechanics, and mechanotransduction. Circ Res 102, 1307–1318, doi:10.1161/CIRCRESAHA.108.173989 (2008).

6 Zhu, J. et al. The effects of measurement parameters on the cancerous cell nucleus characterisation by atomic force microscopy in vitro. J Microsc 287, 3–18, doi:10.1111/jmi.13104 (2022).

7 Wang, K., Qin, Y. & Chen, Y. In situ AFM detection of the stiffness of the in situ exposed cell nucleus. Biochim Biophys Acta Mol Cell Res 1868, 118985, doi:10.1016/j.bbamcr.2021.118985 (2021).

8 Liu, H. et al. In situ mechanical characterization of the cell nucleus by atomic force microscopy. ACS Nano 8, 3821–3828, doi:10.1021/nn500553z (2014).

9 Wilhelm, C. Out-of-Equilibrium Microrheology inside Living Cells. Pys. Rev. Lett. 101, 028101 (2008).

10 Bausch, A., Möller, W. & Sackmann, E. Measurement of Local Viscoelasticity and Forces in Living Cells by Magnetic Tweezers. Biophys J 76, 573–579 (1999).

11 Wu, P.-H. et al. A comparison of methods to assess cell mechanical properties. Nature Methods 15, 491-+, doi:10.1038/s41592-018-0015-1 (2018).

12 Patteson, A. E. et al. Loss of Vimentin Enhances Cell Motility through Small Confining Spaces. Small 15, e1903180, doi:10.1002/smll.201903180 (2019).

13 Loneker, A. E. et al. Lipid droplets are intracellular mechanical stressors that impair hepatocyte function. Proc Natl Acad Sci U S A 120, e2216811120, doi:10.1073/pnas.2216811120 (2023).

14 Ivanovska, I. L., Tobin, M. P., Bai, T., Dooling, L. J. & Discher, D. E. Small lipid droplets are rigid enough to indent a nucleus, dilute the lamina, and cause rupture. J Cell Biol 222, doi:10.1083/jcb.202208123 (2023).

15 Goldman, R. D., Pollack, R. & Hopkins, N. H. Preservation of normal behavior by enucleated cells in culture. Proc Natl Acad Sci U S A 70, 750–754, doi:10.1073/pnas.70.3.750 (1973).

16 Colonna, A. & Kerr, S. J. The nucleus as the site of tRNA methylation. J Cell Physiol 103, 29–33, doi:10.1002/jcp.1041030105 (1980).

17 Ohara, J., Sekiguchi, T. & Watanabe, T. Purification of cytoplasts (enucleated cells) and karyoplasts (nuclei) from mouse splenic lymphocytes with cytochalasin-B. J Immunol Methods 45, 239–248, doi:10.1016/0022-1759(81)90302-1 (1981).

18 Patteson, A. E. et al. Vimentin protects cells against nuclear rupture and DNA damage during migration. Journal of Cell Biology 218, 4079–4092, doi:10.1083/jcb.201902046 (2019).

19 Wu, Y., Pegoraro, A. M., Weitz, D., Janmey, P. M. & Sun, S. The correlation between cell and nucleus size is explained by an eukaryotic cell growth model. Plos Computational Biology 18, doi:10.1371/journal.pcbi.1009400 (2022).

20 Li, Z. et al. Membrane tether formation from outer hair cells with optical tweezers. Biophys J 82, 1386–1395, doi:10.1016/S0006-3495(02)75493-3 (2002).

21 Pontes, B. et al. Cell cytoskeleton and tether extraction. Biophys J 101, 43–52, doi:10.1016/j.bpj.2011.05.044 (2011).

22 Davidson, P. & Lammerding, J. Broken nuclei – lamins, nuclear mechanics and disease. Trends in Cell Biology 24, 247–256 (2014).

23 Pfeifer, C. R. et al. Gaussian curvature dilutes the nuclear lamina, favoring nuclear rupture, especially at high strain rate. Nucleus 13, 129–143, doi:10.1080/19491034.2022.2045726 (2022).

24 Xia, Y. et al. Nuclear rupture at sites of high curvature compromises retention of DNA repair factors. J Cell Biol 217, 3796–3808, doi:10.1083/jcb.201711161 (2018).

25 Sivaramakrishnan, S., DeGiulio, J. V., Lorand, L., Goldman, R. D. & Ridge, K. M. Micromechanical properties of keratin intermediate filament networks. Proc Natl Acad Sci U S A 105, 889–894, doi:10.1073/pnas.0710728105 (2008).

26 Bieluszewski, T., Prakash, S., Roule, T. & Wagner, D. The Role and Activity of SWI/SNF Chromatin Remodelers. Annu Rev Plant Biol 74, 139–163, doi:10.1146/annurev-arplant-102820-093218 (2023).

27 Pogoda, K. et al. Unique Role of Vimentin Networks in Compression Stiffening of Cells and Protection of Nuclei from Compressive Stress. Nano Lett 22, 4725–4732, doi:10.1021/acs.nanolett.2c00736 (2022).

28 Guo, M. et al. Cell volume change through water efflux impacts cell stiffness and stem cell fate. Proc Natl Acad Sci U S A 114, E8618–E8627, doi:10.1073/pnas.1705179114 (2017).

29 Wu, Y., Pegoraro, A. F., Weitz, D. A., Janmey, P. & Sun, S. X. The correlation between cell and nucleus size is explained by an eukaryotic cell growth model. PLoS Comput Biol 18, e1009400, doi:10.1371/journal.pcbi.1009400 (2022).

30 Deviri, D. & Safran, S. A. Balance of osmotic pressures determines the nuclear-to-cytoplasmic volume ratio of the cell. Proc Natl Acad Sci U S A 119, e2118301119, doi:10.1073/pnas.2118301119 (2022).

31 Lemiere, J., Real-Calderon, P., Holt, L. J., Fai, T. G. & Chang, F. Control of nuclear size by osmotic forces in Schizosaccharomyces pombe. Elife 11, doi:10.7554/eLife.76075 (2022).

32 Goswami, R. et al. Mechanical Shielding in Plant Nuclei. Curr Biol 30, 2013–2025 e2013, doi:10.1016/j.cub.2020.03.059 (2020).

33 Cruz, K., Wang, Y.-H., Oake, S. A. & Janmey, P. A. Polyelectrolyte Gels Formed by Filamentous Biopolymers: Dependence of Crosslinking Efficiency on the Chemical Softness of Divalent Cations. Gels 7, 41 (2021).

34 Lammerding, J. et al. Lamins A and C but not lamin B1 regulate nuclear mechanics. J Biol Chem 281, 25768–25780, doi:10.1074/jbc.M513511200 (2006).

35 Stephens, A. D., Banigan, E. J., Adam, S. A., Goldman, R. D. & Marko, J. F. Chromatin and lamin A determine two different mechanical response regimes of the cell nucleus. Mol Biol Cell 28, 1984–1996, doi:10.1091/mbc.E16-09-0653 (2017).

36 Pajerowski, J. D., Dahl, K. N., Zhong, F. L., Sammak, P. J. & Discher, D. E. Physical plasticity of the nucleus in stem cell differentiation. Proc Natl Acad Sci U S A 104, 15619–15624, doi:10.1073/pnas.0702576104 (2007).

37 Denais, C. M. et al. Nuclear envelope rupture and repair during cancer cell migration. Science 352, 353–358, doi:10.1126/science.aad7297 (2016).

38 Keizer, V. I. P. et al. Live-cell micromanipulation of a genomic locus reveals interphase chromatin mechanics. Science 377, 489-+, doi:ARTN eabi9810 10.1126/science.abi9810 (2022).

39 Sirinakis, G. et al. The RSC chromatin remodelling ATPase translocates DNA with high force and small step size. EMBO J 30, 2364–2372, doi:10.1038/emboj.2011.141 (2011).

40 Zhang, Y. et al. DNA translocation and loop formation mechanism of chromatin remodeling by SWI/SNF and RSC. Mol Cell 24, 559–568, doi:10.1016/j.molcel.2006.10.025 (2006).

41 Chang, L. et al. The SWI/SNF complex is a mechanoregulated inhibitor of YAP and TAZ. Nature 563, 265–269, doi:10.1038/s41586-018-0658-1 (2018).

42 Rando, O. J., Zhao, K., Janmey, P. & Crabtree, G. R. Phosphatidylinositol-dependent actin filament binding by the SWI/SNF-like BAF chromatin remodeling complex. Proc Natl Acad Sci U S A 99, 2824–2829, doi:10.1073/pnas.032662899 (2002).

43 Louvet, E., Yoshida, A., Kumeta, M. & Takeyasu, K. Probing the stiffness of isolated nucleoli by atomic force microscopy. Histochem Cell Biol 141, 365–381, doi:10.1007/s00418-013-1167-9 (2014).

44 Mandal, K. et al. Soft Hyaluronic Gels Promote Cell Spreading, Stress Fibers, Focal Adhesion, and Membrane Tension by Phosphoinositide Signaling, Not Traction Force. ACS Nano 13, 203–214, doi:10.1021/acsnano.8b05286 (2019).

45 Murrell, M. P. et al. Liposome adhesion generates traction stress. Nat Phys 10, 163–169, doi:10.1038/Nphys2855 (2014).

46 Goldman, R. D., Pollack, R., Chang, C. M. & Bushnell, A. Properties of enucleated cells. III. Changes in cytoplasmic architecture of enucleated BHK21 cells following trypsinization and replating. Exp Cell Res 93, 175–183, doi:10.1016/0014-4827(75)90437-1 (1975).

47 Rodionov, V., Nadezhdina, E., Peloquin, J. & Borisy, G. Digital fluorescence microscopy of cell cytoplasts with and without the centrosome. Methods Cell Biol 67, 43–51, doi:10.1016/s0091-679x(01)67004-3 (2001).

48 McMullen, A., Munoz Basagoiti, M., Zeravcic, Z. & Brujic, J. Self-assembly of emulsion droplets through programmable folding. Nature 610, 502–506, doi:10.1038/s41586-022-05198-8 (2022).

49 McMullen, A., Hilgenfeldt, S. & Brujic, J. DNA self-organization controls valence in programmable colloid design. Proc Natl Acad Sci U S A 118, doi:10.1073/pnas.2112604118 (2021).

50 Zhang, Y. et al. Sequential self-assembly of DNA functionalized droplets. Nat Commun 8, 21, doi:10.1038/s41467-017-00070-0 (2017).

51 Brettschneider, J. et al. Rapid Lipid Droplet Isolation Protocol Using a Well-established Organelle Isolation Kit. J Vis Exp, doi:10.3791/59290 (2019).

52 Lherbette, M. et al. Atomic Force Microscopy micro-rheology reveals large structural inhomogeneities in single cell-nuclei. Sci Rep 7, 8116, doi:10.1038/s41598-017-08517-6 (2017).

53 Klymenko, O., Wiltowska-Zuber, J., Lekka, M. & Kwiatek, W. Energy dissipation in the AFM elasticity measurements. Acta physica polonica a 115, 548–551 (2009).

