## Supplemental Figure 1 for "Metabolically intact nuclei are fluidized by the activity of the chromatin remodeling motor BRG1"

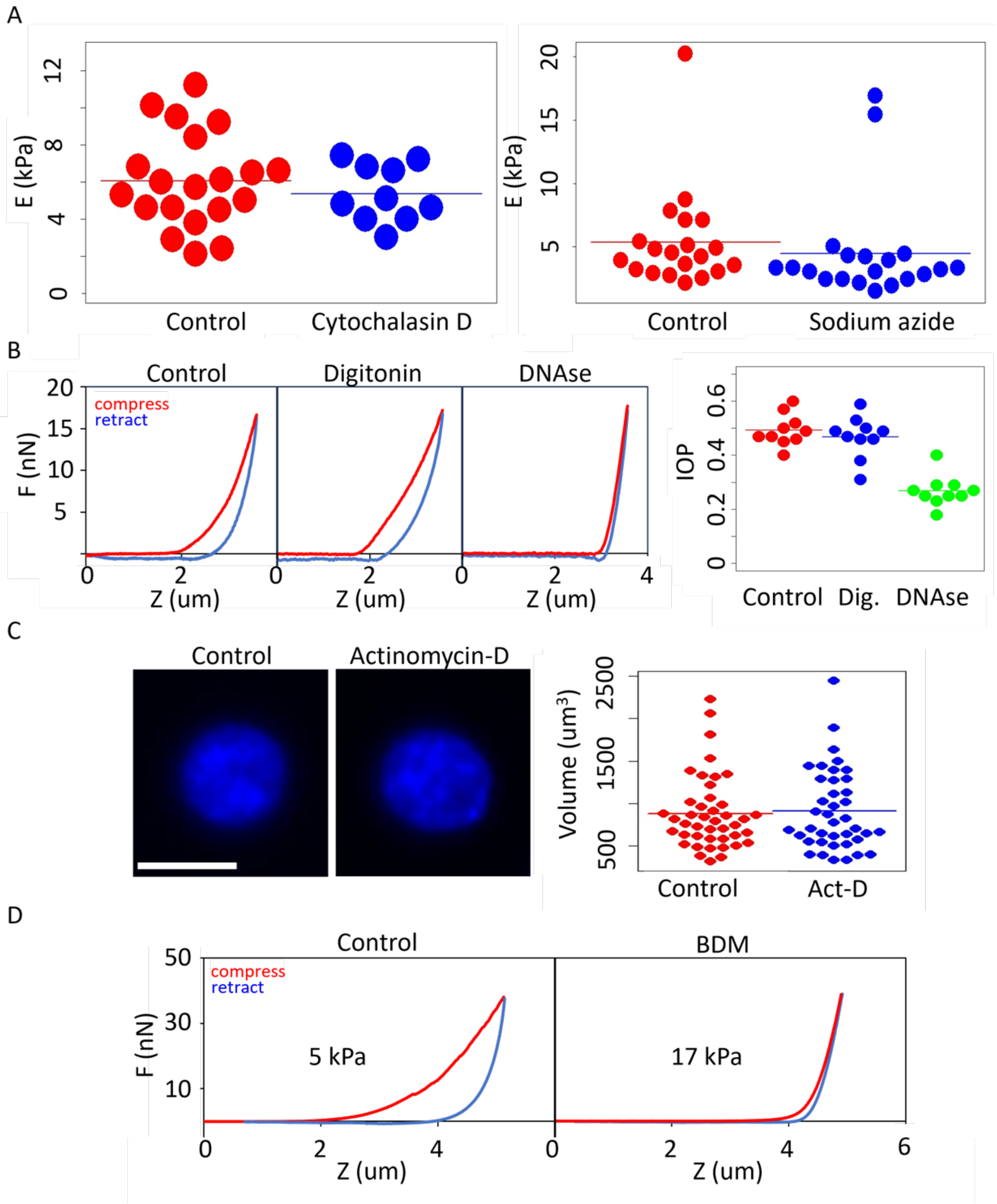


Supplement Figure 1. (A) Scatter plots of karyoplast Young's moduli after addition of 2 ug/ml cytochalasin-D or 2 mM sodium azide for 1 hour. (B) Effects of DNase I on stiffness and dissipation in karyoplasts. Force-recovery plots of control karyoplasts and after treatment with digitonin or digitonin + DNase I. Scatter plot of plasticity index. (C) Effects of RNA polymerase inhibition on karyoplast morphology. Hoechst-stained nuclei and karyoplast volumes before and after treatment with actinomycin-D. Scale bar = 10 µm. (D) Effect of BDM on karyoplast rheology
